# Interrogation of human hematopoiesis at single-cell and single-variant resolution

**DOI:** 10.1101/255224

**Authors:** Caleb A. Lareau, Jacob C. Ulirsch, Erik L. Bao, Leif S. Ludwig, Michael H. Guo, Christian Benner, Ansuman T. Satpathy, Rany Salem, Joel N. Hirschhorn, Hilary K. Finucane, Martin J. Aryee, Jason D. Buenrostro, Vijay G. Sankaran

**Affiliations:** Broad Institute of MIT and Harvard, Cambridge, MA 02142, USA.; Department of Biostatistics, Harvard T.H. Chan School of Public Health.; Department of Pathology, Massachusetts General Hospital.; Division of Hematology/Oncology, The Manton Center for Orphan Disease Research, Boston Children’s Hospital, Harvard Medical School, Boston, MA 02115, USA.; Department of Pediatric Oncology, Dana-Farber Cancer Institute, Harvard Medical School, Boston, MA 02115, USA.; Program in Biological and Biomedical Sciences, Harvard Medical School, Boston, MA 02115, USA.; Harvard-MIT Health Sciences and Technology, Harvard Medical School, Boston, MA 02115, USA.; Division of Endocrinology, Boston Children’s Hospital, Harvard Medical School, Boston, MA 02115.; Department of Genetics, Harvard Medical School, Boston, MA 02115.; Center for Basic and Translational Obesity Research, Boston Children’s Hospital, Boston, MA 02115.; Institute for Molecular Medicine Finland, University of Helsinki, 00014 Helsinki, Finland.; Department of Public Health, University of Helsinki, 00014 Helsinki, Finland.; Department of Pathology, Stanford University School of Medicine, Stanford, CA, 94305.; Harvard Society of Fellows, Harvard University, Cambridge, MA 02138, USA.

## Abstract

Incomplete annotation of cell-to-cell state variance and widespread linkage disequilibrium in the human genome represent significant challenges to elucidating mechanisms of trait-associated genetic variation. Here, using data from the UK Biobank, we perform genetic fine-mapping for 16 blood cell traits to quantify posterior probabilities of association while allowing for multiple independent signals per region. We observe an enrichment of fine-mapped variants in accessible chromatin of lineage-committed hematopoietic progenitor cells. Further, we develop a novel analytic framework that identifies “core gene” cell type enrichments and show that this approach uniquely resolves relevant cell types within closely related populations. Applying our approach to single cell chromatin accessibility data, we discover significant heterogeneity within classically defined multipotential progenitor populations. Finally, using several lines of empirical evidence, we identify relevant cell types, predict target genes, and propose putative causal mechanisms for fine-mapped variants. In total, our study provides an analytic framework for single-variant and single-cell analyses to elucidate putative causal variants and cell types from GWAS and high-resolution epigenomic assays.

## Introduction

Hematopoiesis is a well-characterized paradigm of cellular differentiation that is highly coordinated and regulated to ensure balanced proportions of mature blood cells.^1^ Despite our sophisticated understanding gained primarily from model organisms, many aspects of this process remain poorly understood in humans. At the population level, there is a wide spectrum of phenotypic variation in commonly measured blood cell traits, such as hemoglobin levels and specific blood cell counts, which can manifest as various diseases at extreme ends of the spectrum.^2^ Identifying genetic variants that drive these differences in blood cell traits in human populations may reveal important regulatory mechanisms and genes critical for blood cell production and diseases.

To these ends, genome-wide association studies (GWAS) have identified thousands of genomic loci linked to complex phenotypes such as blood cell traits,^3^ but a major challenge has been the identification of causal genetic variants and relevant cell types underlying the observed phenotypic associations.^4^ In particular, linkage disequilibrium (LD) has confounded the precise identification of functional variant(s). In an effort to better identify casual variants, several statistical approaches have been developed. The first, termed *genetic fine-mapping*, attempts to resolve trait-associated loci to likely causal variants by modeling LD structure and the strength of associations. In practice, a major limitation has been that most of these methods assume exactly one causal variant per locus,^5,6^ despite strong evidence that a substantial number of these loci contain multiple independent associations.^7–10^ The second type of approach instead focuses on identifying functional tissue enrichments. It has now been well established that ~80–90% of associated loci do not tag coding variants and that ~40–80% of the narrow-sense heritability of many complex traits can be resolved to genomic regulatory regions.^11,12^ Given this observation, tissue-specific measurements of regulatory element activity are often overlapped with significant association loci (e.g. *epigenomic fine-mapping*) or investigated genome-wide (e.g. partitioned heritability) in order to identify variants and cell types most likely to underlie the measured phenotypic trait.^11,13^ These enrichment methods have revealed causal tissues for diseases, such as pancreatic islets in diabetes^13^ and central nervous system cells in schizophrenia,^11^ but these methods are only beginning to be applied to highly related traits and cell types within single systems, such as the hematopoietic hierarchy.

Recently, advances in single cell genomic approaches have allowed us to define discrete cell types in complex systems and have been used to further revise our understanding of the hematopoietic hierarchy. For example, single-cell transcriptomic and chromatin accessibility studies have identified previously undescribed progenitor populations, subdivided classically defined populations into distinct cell types, and revealed that lineage commitment occurs in earlier progenitors than previously appreciated.^14–19^ Nevertheless, the functional consequences of human genetic variation in relation to these refined models of cell fate commitment in hematopoiesis, as reflected in peripheral blood measurements, have not yet been explored.

Given that blood cell counts provide direct phenotypic readouts of hematopoietic lineage commitment and differentiation, we reasoned that we could use variants significantly associated with these phenotypes to dissect hematopoiesis in an unbiased fashion. In this study, we performed a GWAS on ~113,000 “White British” individuals from the UK Biobank, similar to a previously described study.^3^ Leveraging this large sample size and precise population LD information, we fine-mapped multiple causal variants in hundreds of individual regions. Furthermore, we describe and validate a method (g-chromVAR) to discriminate between closely related cell types in an effort to identify relevant stages of hematopoiesis that are affected by these common genetic variants. Here, rather than utilizing summary statistics from all variants (for an “omnigenic” trait^4^), we demonstrate specificity using our “core” subset of fine-mapped variants. We show that this approach can be used to score chromatin accessibility from individual cells for GWAS enrichment, and we use these enrichments to interrogate both the timing and variability of lineage commitment in 2,034 single cells. Finally, we provide a community resource comprised of our fine-mapping results combined with several layers of functional evidence.

## Results

### Fine-mapping pinpoints hundreds of likely causal variants

We performed genome-wide association studies (GWASs) for 16 blood cell traits on the UK Biobank (UKBB), which consists of 113,000 individuals of predominantly European descent. These blood cell traits represent several important and distinct hematopoietic lineages, including red blood cells (RBCs), platelets, lymphocytes (T and B cells), neutrophils, monocytes, eosinophils, and basophils (**Fig. 1A**). Using LD-score regression (LDSR),^20^ we found that common variants of minor allele frequency (MAF) > 1% captured moderately high narrow sense heritability 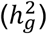 in most traits, with an average of 15.4%, ranging from 4% (basophil count) to 32% (mean platelet volume) (**Fig. S1A**). In general, traits from the same lineage, such as RBC count and hemoglobin, had high genetic correlations (R^2^ = 0.91, Bonferroni p < 0.001) (**Fig. S1C**). However, several traits from distinct lineages also had significant genetic correlations, such as platelet count and lymphocyte count (R^2^ = 0.29, Bonferroni p < 0.001), underscoring the presence of genetic pleiotropy among the hematopoietic traits (**Fig. S1B**). These analyses confirmed that common genetic variants contribute substantially to blood cell trait heritability and that regulation of hematopoiesis could potentially occur across various stages of hematopoiesis.

**Figure 1.**
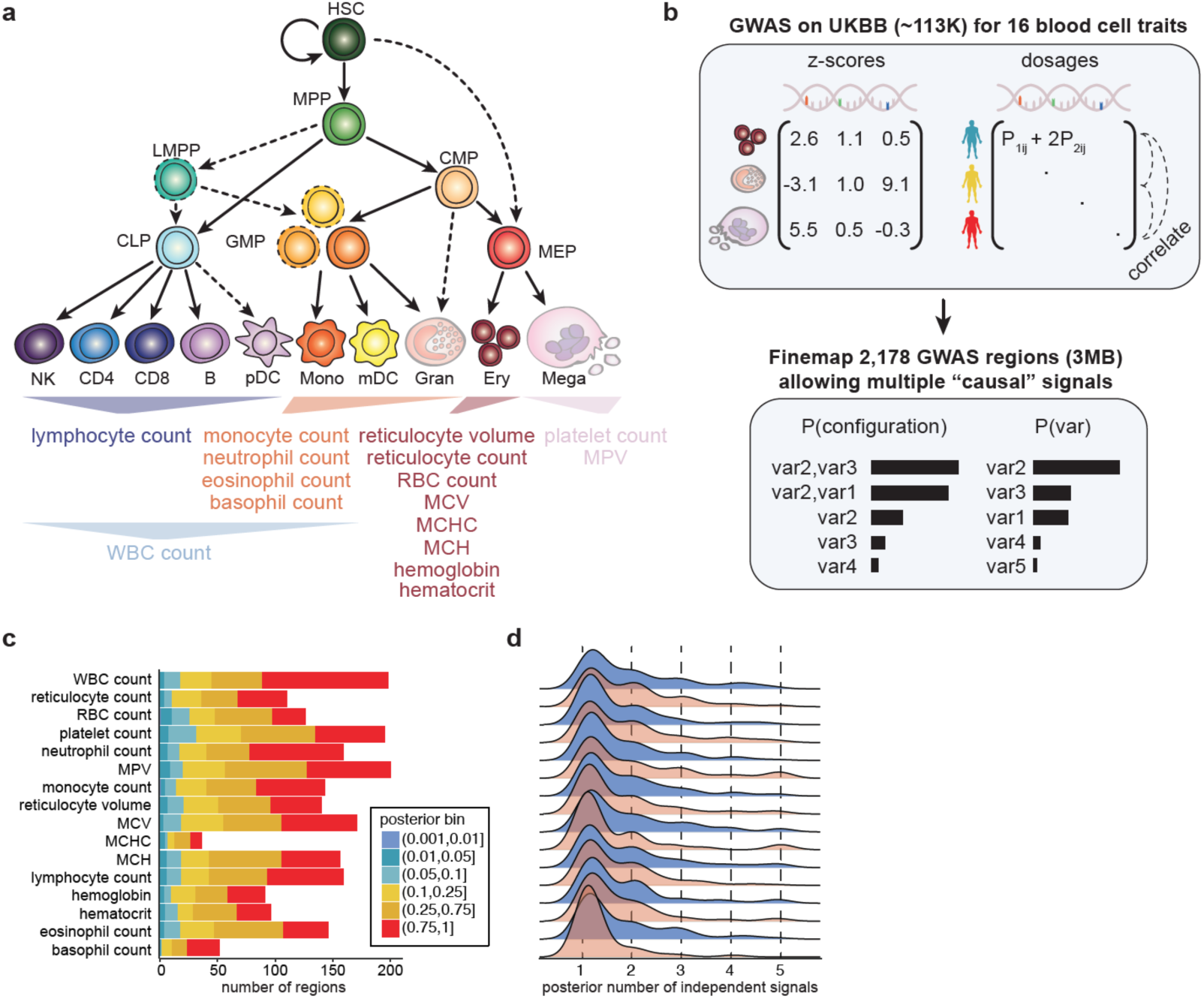
Overview of hematopoiesis, UKBB GWAS, and fine-mapping, (**a**) Schematic of the human hematopoietic hierarchy showing the primary cell types analyzed in this work. ATAC-seq and RNA-seq were collected for all sorted cell types except granulocytes (ATAC-seq, RNA-seq) and megakaryocytes (RNA-seq). Colors used in this schematic are consistent throughout all figures. Mono, monocyte; gran, granulocyte; ery, erythroid; mega, megakaryocyte; CD4, CD4+ T cell; CD8, CD8+ T cell; B, B cell; NK, natural killer cell; mDC, myeloid dendritic cell; pDC, plasmacytoid dendritic cell. The 16 terminal blood traits that were genetically fine-mapped are shown below the hierarchy. (**b**) Schematic of UKBB GWAS and fine-mapping approach. Briefly, blood traits from ~113K individuals were fine-mapped allowing for multiple causal variants and using imputed genotype dosages as reference LD. (**c**) Number of fine-mapped 3MB regions for each trait with the best posterior probability for a variant being causal indicated. (**d**) Kernel density of the expected number of causal variants for each region.

To begin to dissect the nature and timing of the effects of common genetic variation during hematopoiesis, we performed genetic fine-mapping to identify high confidence variants. Traditional fine-mapping approaches operate under the assumption that there is only one causal variant per locus and are either agnostic to LD or use small whole genome sequencing (WGS) reference panels from “matched” populations, which have been shown to be inaccurate when scaled to large sample sizes.^21^ To overcome these limitations, we first identified 2,178 regions (each 3 MB in size) across the genome, excluding the HLA region due to its notoriously complex LD structure, that were associated with at least one blood cell trait (minimum p-value < 5 × 10^−8^; see Online Methods). For each region, we calculated LD directly from the imputed genotype probabilities (dosages) using all individuals included in our blood cell trait GWASs, rather than from a hard-called reference panel (**Fig. 1B**). Finally, we performed Bayesian fine-mapping on all common variants (MAF > 0.1%) with satisfactory imputation (INFO > 0.6).^22^

Across these 2,178 regions, our method identified 36,919 variants with >1% posterior probability (PP) of being causal for a trait association (**Table S1, S2**), and 1,110 regions (51%) contained at least one variant with >50% PP (**Fig. 1C**), providing strong evidence that our approach was successful in pinpointing causal variants. One advantage of our approach is that it allows for the identification of multiple independent signals per associated locus: starting with a conservative prior, the posterior expected number of independent causal variants was >2 for 36% of regions and >3 for 13% of regions (**Fig. 1D**). Moreover, the fine-mapped variants captured a significant proportion of narrow sense heritability (trait average of 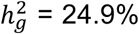 for PP > 1%) (**Fig. S1A**), suggesting that our approach identifies a core subset of variants that explains a large amount of phenotypic variation.

As an example of a locus with multiple independent signals, we investigated the *CCND3* locus in which we previously used trans-ethnic fine-mapping and functional assays to identify a causal variant and associated gene leading to variation in RBC count.^23^ Our approach correctly identified the known causal variant (rs9349205) as the primary association, as well as ~4 additional independent signals in this locus, including a secondary imputed variant (rs112233623) associated with decreased RBC count (**Fig. 2A-C**). As further validation of our approach, stepwise conditional analysis also identified 4 independent significant signals and the same secondary variant (rs112233623) in this locus (**Fig. 2B**). Importantly, this secondary variant was not identified by fine-mapping if we instead used LD estimated from either the UK10K WGS reference panel (**Fig. 2D**) or hard-called variants from the UKBB population (**Fig. S2**), highlighting the value of calculating LD using imputed genotype dosages from the GWAS population itself (note that BOLT-LMM uses these dosages when determining associations). Interestingly, rs112233623 is only 161 bases from the original causal variant (rs9349205), and both lie within an erythroid-specific nucleosome depleted region (NDR) (**Fig. 2E**). We investigated the potential synergy between these two variants using luciferase reporter assays and found that each variant affected enhancer activity independently (p = 0.00178 for rs9349205; p = 2.86e-06 for rs112233623) with minor allele effects in opposing directions, consistent with the genetic directionality and our previous functional work^23^ on rs9349205 and RBC count (**Fig. 2F**). We further verified independent effects for two fine-mapped variants (rs49950 and rs12005199; PP > 0.999; 123 base pairs apart) associated with platelet count within a single NDR ~20kb upstream of *AK3*, a gene whose zebrafish homolog is essential for platelet (thrombocyte) formation (**Fig. S3**).^24^ Notably, we did not observe a significant multiplicative (epistatic) effect at either locus (p = 0.129 for *CCND3*; p = 0.238 for *AK3*). Thus, our statistical and genetic characterization of the *CCND3* and *AK3* loci show that multiple independent causal variants can occur not only in the same LD-block but in the *same regulatory element*. These results indicate that other fine-mapping methodologies that assume one causal variant per locus likely miss true independent effects.

**Figure 2.**
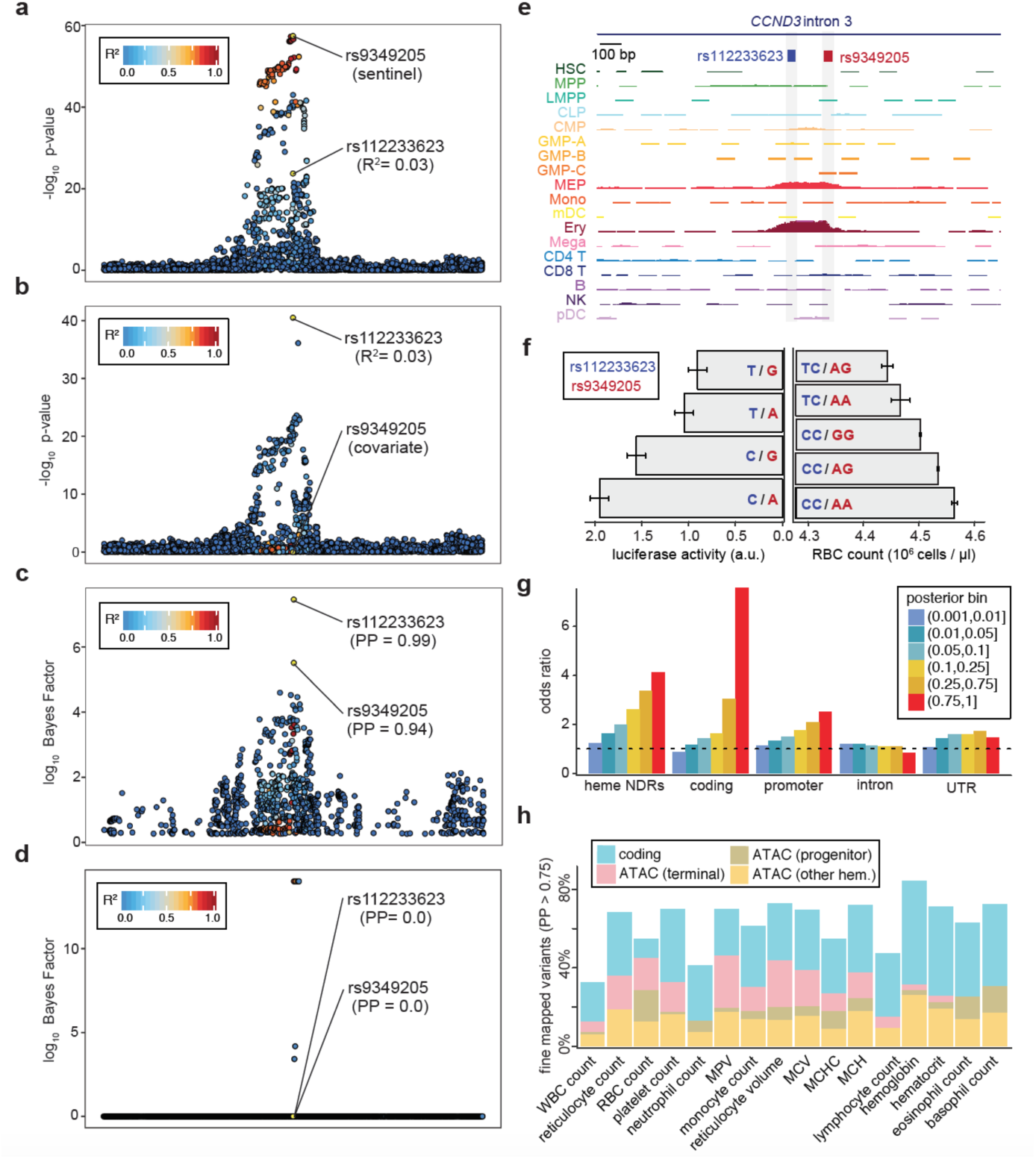
Characterization and validation of *CCND3* intronic region with multiple causal variants. Regional association plots for RBC count (**a**) from the initial GWAS and (**b**) after conditioning on the sentinel variant rs9349205. (**c**) Fine-mapping identifies two putative causal variants (rs9349205, PP=0.94; rs112233623, PP=0.99), but (**d**) not when performed using an LD reference panel of 3,677 WGS individuals from the UK10K study. (**e**) Both putative causal variants (161 base pairs apart) lie within the same erythroid-specific nucleosome-depleted region (NDR). (**f**) Luciferase reporter assays for four haplotypes (left) corroborate independent additive effects of rs9349205 (red) and rs112233623 (blue) on RBC count (right). (**g**) Local-shifting enrichments of fine-mapped variants across all traits for varying posterior probability bins. (**h**) Annotation of PP >0.75 variants per trait indicating overlap in coding and NDR regions across 18 ATAC-seq samples.

In order to further validate our fine-mapping approach, we investigated the overlap of our fine-mapped variant sets (binned by PP) with several key genomic annotations previously shown to be enriched for GWAS signals.^11,12^ To generate a null distribution, we locally shifted annotations within a 3 MB window, similar to the method implemented in GoShifter.^25^ We observed minimal enrichment for intronic and untranslated regions of genes across all bins, but found strong, focal, and stepwise enrichments across higher PP bins for hematopoietic NDRs, promoters, and coding regions (OR=4.2, 2.9, and 8.5 for PP > 75% respectively), consistent with the enrichments reported in these annotations for other complex traits (**Fig. 2G**).^11,12,25^ When we excluded all variants with high correlation (R^2^ > 0.8) to the sentinel variants, we found similarly strong enrichments in hematopoietic NDRs (**Fig. S2**), further suggesting that the fine-mapped secondary and tertiary signals are likely to be functional. Overall, an average of 62% of the most fine-mapped variants (PP > 75%) were either protein coding variants or resided within hematopoietic enhancers (**Fig. 2H**). Notably, the large proportion of fine-mapped coding variants can be attributed to the dense genotyping and large sample size from the UKBB that was well-powered to detect associations for lower frequency variants.

### Development of g-chromVAR, a novel method to measure fine-mapped GWAS set enrichment in continuous annotations

We next set out to determine the exact stages of human hematopoiesis at which *regulatory* genetic variation underlying each blood cell trait was most likely acting. In this well-defined system of cellular differentiation, a strong association in a specific cell population would provide evidence that genetic variation is regulating the transcription of genes important for a particular phenotype (e.g. RBC count, lymphocyte count) in that specific cell type. Although recent methods^11,25^ have been developed to calculate enrichment of genetic variation with genomic annotations, a method which takes into account both **(1)** the strength and specificity of the genomic annotation and **(2)** the probability of variant causality, accounting for LD structure, is needed to resolve associations within the stepwise hierarchies that define hematopoiesis. To these ends, we developed a new approach called genetic-chromVAR (**g-chromVAR**), a generalization of the recently described chromVAR method^26^ to measure the enrichment of regulatory variants in each cell state using uncertainties in fine-mapped genetic variants and quantitative measurements of regulatory activity (**Fig. 3A**, see online methods for additional details). Briefly, this method weights chromatin features by variant posterior probabilities and computes the enrichment for each cell type versus an empirical background matched for GC content and feature intensity (g-chromVAR is thus intuitively a *competitive* method across cell types). We show that g-chromVAR is generally robust to variant posterior probability thresholds and numbers of background peaks (**Fig. S4B,C**) and captures true enrichments in a simulated setting (**Fig. S5**).

**Figure 3.**
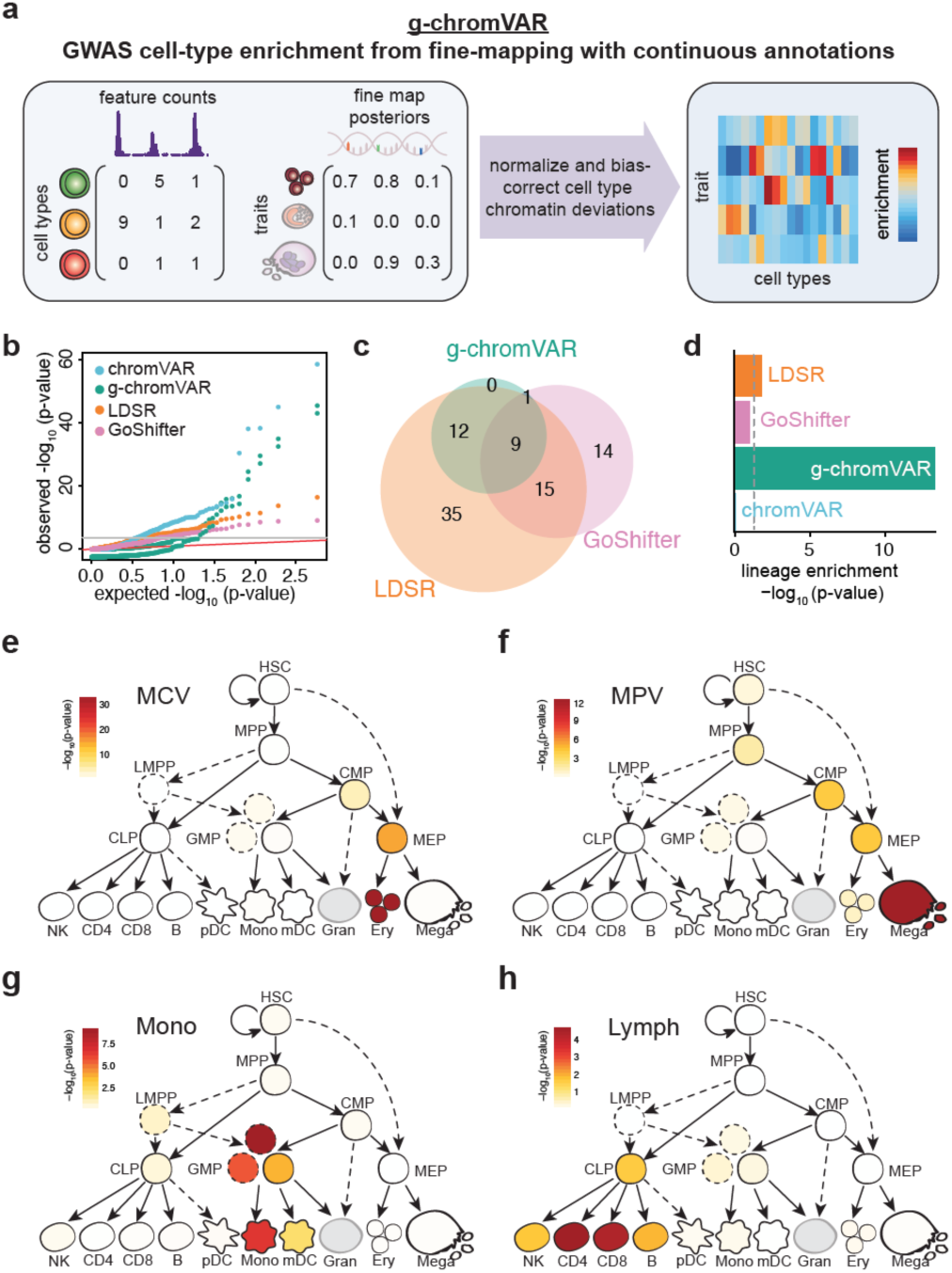
Overview of g-chromVAR and application to hematopoietic cell types. (**a**) Schematic showing inputs for quantitative epigenomic data for each cell type (**X**) and the matrix of fine-mapped variant posterior probabilities for GWAS traits (**G**). Results from the application of g-chromVAR and three similar methods to 16 blood cell traits for 18 hematopoietic cell types are shown in **b**-**d**. (**b**) Quantile-quantile representation of the p-values from each method. (**c**) Overlap between methods for Bonferroni-corrected trait enrichments. (**d**) Lineage-enrichment of all trait-pairs for each method. A Mann-Whitney rank-sum test was used to evaluate the relative enrichment of lineage-specific trait-cell type pairs (true positives). Representative enrichments for 4 traits using g-chromVAR: (**e**) mean corpuscular volume; (**f**) mean platelet volume; (**g**) monocyte count; (**h**) lymphocyte count.

We applied g-chromVAR to each of the 16 blood cell traits and 18 chromatin accessibility profiles (ATAC-seq) of hematopoietic progenitor and terminal populations primarily sorted from the bone marrow or blood of multiple healthy donors (**Fig. 1A**, **Table S3**).^27,28^ We compared g-chromVAR to two state-of-the-art methods: LDSR,^11^ which calculates the enrichment for genome-wide heritability using binary annotations after accounting for LD and overlapping annotations, and GoShifter,^25^ which calculates the enrichment of tight LD blocks containing sentinel GWAS single nucleotide variants for binary annotations. Using a Bonferroni correction, g-chromVAR identified 22 trait-tissue enrichments, LDSR identified 71, GoShifter identified 39, and chromVAR identified 79 (**Fig. 3C**).

In order to compare the performance of these enrichment tools, we leveraged our knowledge of the hematopoietic system and devised a *lineage specificity test*. For any measured cell trait, we identified all possible upstream progenitor populations that could be passed through before terminal differentiation (**Fig. 1A**). For example, the differentiation of an RBC is thought to begin at the hematopoietic stem cell (HSC) and progress through multipotent progenitor (MPP), common myeloid progenitor (CMP), and megakaryocyte erythroid progenitor (MEP) before reaching the erythroid progenitor (Ery) stage. The *lineage specificity test* is a nonparametric rank-sum test that measures the relative ranking of *lineage* specific trait-cell type pairs relative to the *non-lineage* specific traits for each of the compared methodologies. Using this metric for specificity, we found that g-chromVAR vastly outperformed all three other methods (**Fig 3D**). Additionally, we found that 21/22 (95%) of g-chromVAR trait-cell type enrichments were supported by LDSR, all of which were *lineage* specific (**Fig. 3C**, **Table S4**). For certain traits such as monocyte count, we found highly similar enrichment patterns for g-chromVAR and LDSR, but *non-lineage* enrichments for chromVAR (**Fig. S6**). For other traits, such as mean reticulocyte volume, g-chromVAR identified only the two most terminally proximal cell types (MEP and Ery) as significantly enriched for the trait, whereas LDSR non-specifically identified 15/18 of the investigated cell types as enriched after Bonferroni correction (**Fig. S6**). We note that we can improve the lineage specificity of LDSR by including all hematopoietic ATAC-seq annotations in the model as covariates, but this results in a loss of statistical power (**Fig S6**).

We further tested g-chromVAR on chromatin accessibility profiles for 53 tissues across the human body^29^ and identified 40 additional trait-cell type enrichments at an FDR of 1%, including 9 traits in CD8 T-cells and 13 total from 3 human embryonic stem cell lines (**Fig. S7A**, **Table S5**). Extending to non-hematopoietic tissues yielded some compelling findings, such as an enrichment of platelet count in a lung fibroblast cell line (IMR90), consistent with a recent study implicating the lung as a key site of platelet production.^30^ However, increasing the number of non-hematopoietic tissues impairs the background model, resulting in false positive enrichments. Thus, we suggest that g-chromVAR is best suited to resolve specific cell types within individual systems containing closely related cells (e.g. hematopoiesis) rather than a breadth of samples across diverse tissues. Finally, in order to test if alternative fine-mapping methods could be used with g-chromVAR, we investigated 39 predominately immune-related disorders that had previously been fine-mapped with PICS^5^ for enrichments with the 18 hematopoietic progenitor populations. We identified 20 trait-cell type enrichments at an FDR of 1% (**Fig. S7B**, **Table S6**), including RBC traits with erythroid precursors, and multiple autoimmune traits with immune subsets, such as multiple sclerosis with CD8+ T, CD4+ T, B, and NK cells; Crohn’s disease with CD4+ T cells; and Kawasaki disease with B cells. Of particular interest was the observation that variants associated with C-reactive protein (CRP) levels, a non-specific biomarker of inflammation, were most strongly enriched for myeloid dendritic cells (mDCs) and other myeloid cell types, further implicating the role of myeloid cells in acute-phase inflammation.^31–33^

Having validated our approach, we investigated cell type enrichments for each of the 16 traits. We found that the most lineage-restricted populations were typically most strongly enriched (**Fig. 3E-H**). For example, RBC count was most strongly enriched in erythroid progenitors (**Fig. 3E**), platelet count was most strongly enriched in megakaryocytes (**Fig. 3F**), and lymphocyte count was most strongly enriched in CD4+ and CD8+ T cells (**Fig. 3H**). In several instances, we observed significant enrichments for traits in earlier progenitor cell types within each lineage, including enrichment for platelet traits in CMPs and enrichment for monocyte traits in a specific subpopulation of GMPs (**Fig. S4A**). Building on several studies that recently demonstrated transcriptomic^14–19^ and chromatin accessibility heterogeneity^27^ within these populations, we next sought to apply g-chromVAR to single-cell ATAC-seq (scATAC-seq) data in order to interrogate the impact of common genetic variation underlying blood cell traits in heterogeneous cell populations.

### GWAS enrichment in single-cell chromatin accessibility data

We applied g-chromVAR to 2,034 single bone-marrow derived hematopoietic progenitor cells profiled using scATAC-seq.^27^ Although our strongest g-chromVAR enrichments for blood traits were in the most lineage restricted precursors which are not represented in this dataset, we reasoned that investigating progenitor populations that *did* have robust enrichment signals, such as CMPs and MEPs, could inform principles of terminal blood cell production at the single cell level and may allow us to infer subpopulations of lineage-biased progenitors that exist within these heterogeneous populations. To these ends, we scored each single cell for GWAS enrichment for the 16 traits using g-chromVAR (**Fig. 4A**). In order to validate our approach, since only a small number of NDRs are measured for each cell (~10,000 detected per cell), we compared GWAS enrichments in the composite profiles of each of the 11 immunophenotypically sorted single cell populations to the enrichments in the bulk profiles and observed that they were highly correlated (r = 0.84) (**Fig. 4B**). As such, we reasoned that aggregating cells by either pseudotime trajectories or clustering of single-cell data could provide new biological insights into the effects of genetic variation on dynamic cell fate decisions in hematopoiesis.

**Figure 4.**
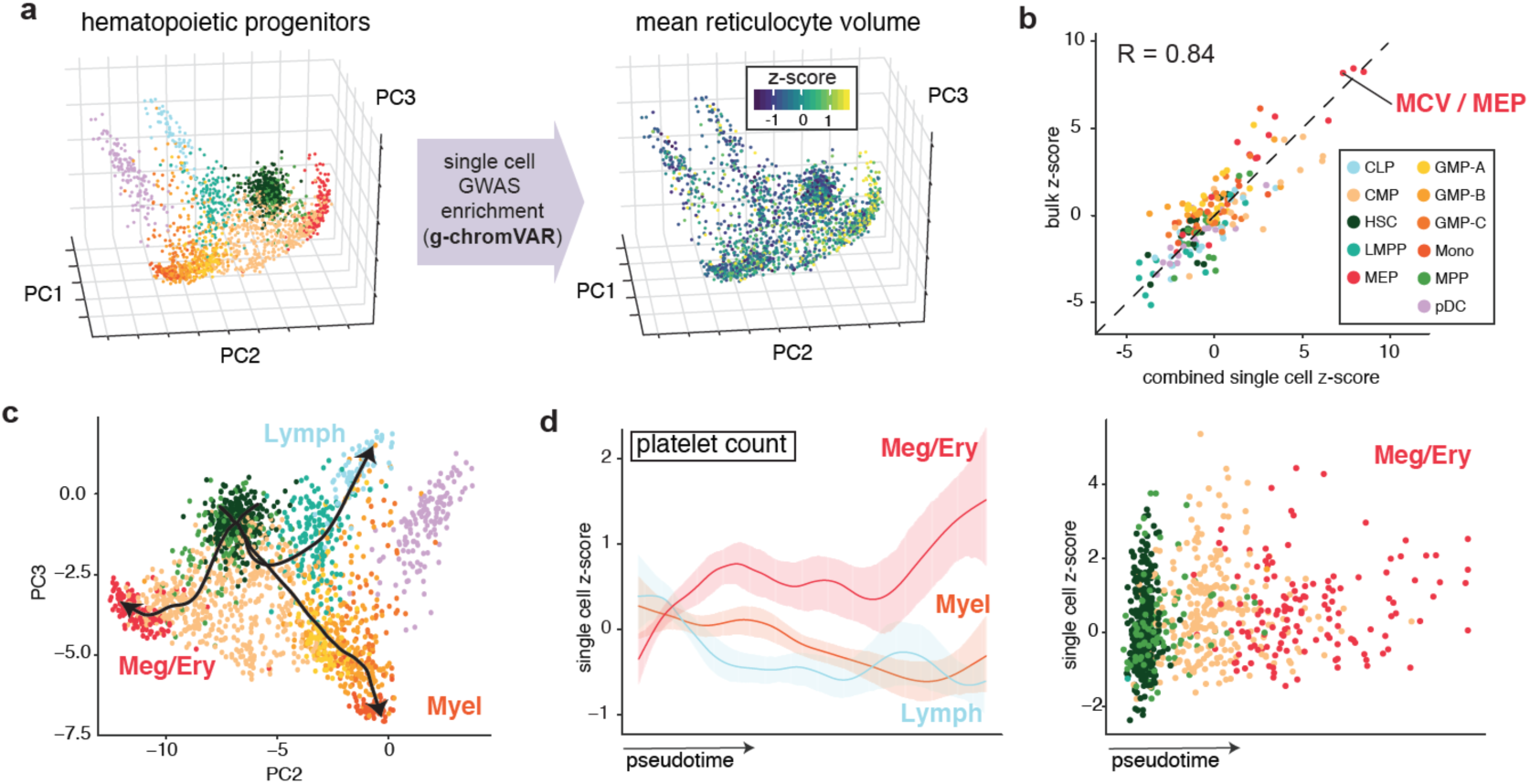
Application of g-chromVAR to single-cell chromatin accessibility data. (**a**) 2,034 hematopoietic single cells projected onto a three-dimensional principal components embedding. Single cells colored by g-chromVAR enrichment scores for mean reticulocyte volume reveal specific regulatory enrichment in the megakaryocyte-erythroid progenitor population. (**b**) Validation of g-chromVAR enrichments using synthetic bulk populations from sums of single cells. Aggregated single-cell g-chromVAR z-scores across all trait-cell type pairs (individual points) strongly correlate (r = 0.84) with bulk population z-scores. (**c**) Representation of inferred pseudotime trajectories of 3 hematopoietic lineages from scATAC-seq data. (**d**) Pseudotime trends of g-chromVAR scores for platelet count across all single cells corroborates regulatory dynamics of megakaryocyte/erythroid differentiation.

In order to model the relatedness and heterogeneity of single cell measurements, we inferred pseudotime trajectories for the megakaryocyte and erythroid lineage (Meg/Ery), the myeloid and monocyte lineage (Myel), and the lymphoid lineage (Lymph) (**Fig. 4C**). We then used local regression to investigate the timing of blood trait GWAS enrichments during lineage commitment. Interestingly, we found that along these trajectories we could reconstruct our observations from bulk data, albeit with finer granularity. For example, we found that platelet count showed enrichment early along Meg/Ery differentiation with a sharp increase at firmly committed MEPs (**Fig. 4D**). This enrichment along the Meg/Ery commitment path coincided with negative enrichments along the alternative Myel and Lymph paths (**Fig. 4D**). This suggests that although the majority of variants act in committed progenitors, a subset of regulatory variants act in multipotential progenitors – in fact, we could identify many distinctly fine-mapped variants that only overlapped with bipotential or multipotential progenitor populations (**Fig. 2H**).

### Regulatory heterogeneity within classically defined hematopoietic populations

Although our developmental trajectories revealed initial insight into the regulation of hematopoietic lineage commitment, GWAS enrichments in each immunophenotypically similar population showed large within-group variation. We thus sought to determine the extent to which this heterogeneity was due to true cell-cell biological differences in regulatory variation. For each of the 16 blood cell traits, we calculated variance in regulatory GWAS enrichment for each of the 11 available hematopoietic progenitor populations. Interestingly, we found that CMP (n=502 cells) and MEP (n=138 cells) populations, in which we identified trait enrichments in the bulk profiles, demonstrated significant heterogeneity in g-chromVAR enrichments for both erythroid and megakaryocyte traits (**Fig. 5A**).

**Figure 5.**
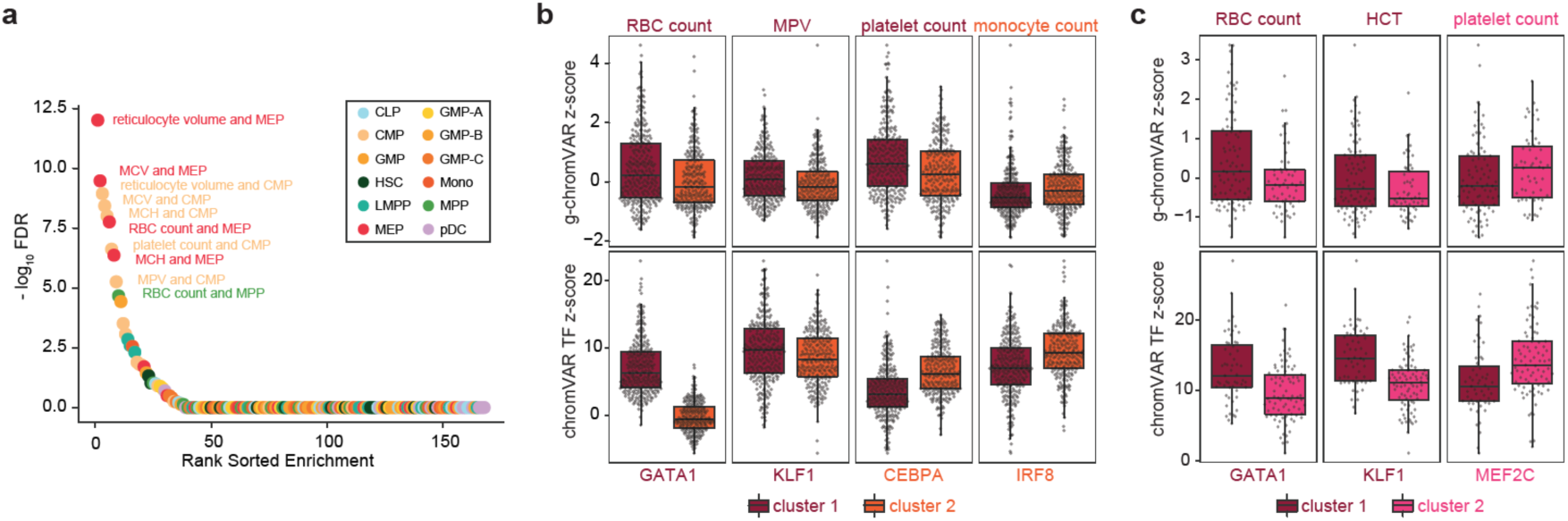
Inference of variable regulatory enrichment in single hematopoietic cells, (**a**) Rank order plot highlighting the trait-cell type pairs with the greatest variance over that of a *χ*^2^ distribution. (**b**) K-medoids partitioning of ATAC-seq counts in CMP cells reveals two subpopulations: monocyte-biased and megakaryocyte/erythroid-biased (RBC count, FDR=0.000128; MPV, FDR=0.000236; platelet count, FDR=1.399e-05; monocyte count, FDR=0.0221). ChromVAR scores for master transcription factors (TFs) of each blood cell type correspond to the lineage-biased clusters (GATA1, FDR=1.759e-82; KLF1, FDR=4.328e-03; CEBPA, FDR=2.575e-16; IRF8, FDR=4.651e-15). (**c**) Similar k-medoids partitioning of MEP cells reveals 2 subpopulations with differential enrichments for megakaryocyte or erythroid phenotypes (RBC count, FDR=0.155; HCT, FDR=0.0398; platelet count, FDR=0.0765), along with consistent differences in chromVAR TF-deviation scores for master TFs of each blood cell type (GATA1, FDR=2.175e-04; KLF1, FDR=4.015e-06; MEF2C, FDR=2.520e-03).

To resolve this heterogeneity, we compared enrichment scores for myeloid and erythroid traits between the CMP and MEP populations. We hypothesized that the CMP population could be subdivided into megakaryocyte/erythrocyte-primed and monocyte-primed subtypes, whereas the MEP population could be further subdivided into erythrocyte-primed and megakaryocyte-primed subtypes. To test this hypothesis, we divided the CMP population into two unsupervised clusters by applying k-medoids to the first 5 principal components of global chromatin accessibility (and naive to g-chromVAR enrichments) (**Fig. S8**). Of note, CMP cluster 1 had significantly higher g-chromVAR enrichments for megakaryocyte and erythrocyte traits, whereas cluster 2 had a significantly higher enrichment for monocyte count (RBC count, FDR=0.000128; MPV, FDR=0.000236; platelet count, FDR=1.399e-05; monocyte count, FDR=0.0221; **Fig. 5B**). Concomitantly, CMP cluster 1 showed higher chromatin accessibility of motifs for erythroid-specific TFs, including GATA1 and KLF1, whereas cluster 2 was enriched for motifs predicted to be bound by the monocyte TFs, CEBPA and IRF8 (GATA1, FDR=1.759e-82; KLF1, FDR=4.328e-03; CEBPA, FDR=2.575e-16; IRF8, FDR=4.651e-15; **Fig. 5B**). Similarly, we clustered the MEP population into two distinct clusters (**Fig. S5**) and found that these clusters were characterized by differences in erythroid and megakaryocyte traits (RBC count, FDR=0.155; HCT, FDR=0.0398; platelet count, FDR=0.0765), as well as chromatin accessibility of motifs for TFs such as MEF2C, a key regulator of megakaryopoiesis (GATA1, FDR=2.175e-04; KLF1, FDR=4.015e-06; MEF2C, FDR=2.520e-03; **Fig. 5C**).^34^ Although additional studies with higher-throughput single cell data are needed to determine whether these differences are due to distinct lineage-biased subpopulations or whether they reflect gradations along a common axis of differentiation, our findings show that heterogeneity in hematopoietic progenitor populations can be linked to putative functional consequences on blood production, reflected in measurable blood cell traits.

### Dissecting the mechanisms of core gene regulation in blood production

Finally, we sought to better understand the precise regulatory mechanisms underlying how fine-mapped genetic variants influence hematopoietic traits. For all 140,750 variants with PP > 0.1%, we combined several lines of functional and predictive evidence to better understand the transcriptional mechanisms and “core genes” involved in blood cell production and to help prioritize variants for experimental validation (**Fig. 6A**). Briefly, we **(1)** calculated TF motif overlap and disruption by each variant for 640 human motifs,^35,36^ **(2)** investigated TF occupancy overlap from 2,115 publicly available ChIP-seq datasets in hematopoietic tissues and cell lines,^37^ **(3)** predicted the effects of each variant on open chromatin in a TF agnostic manner using gkm-SVM,^38^ **(4)** identified empirical target genes by correlating our ATAC-seq with matching RNA-seq data across 17 hematopoietic populations as well as by calculating overlap with recent promoter capture Hi-C,^39^ **(5)** combined our results with whole blood expression quantitative trait loci (eQTLs) and methylation quantitative trait loci (meQTL), and **(6)** performed phenome-wide association studies (pheWAS) for each variant for 642 unique medical ICD-10 diagnoses (**Fig. 6A**). In this effort, we observed that specific fine-mapped variants for blood traits are associated with common diseases such as ischemic heart disease and mineral metabolism disorders (**Fig. 6B**). Furthermore, we verified the reported mechanisms of 9 previously reported causal variants that we were able to fine-map (**Fig. 2**, **Fig. S9**). We provide this compendium of information in an interactive web-based portal as a publicly-accessible resource (**Fig. S10**, see **URLs**).

**Figure 6.**
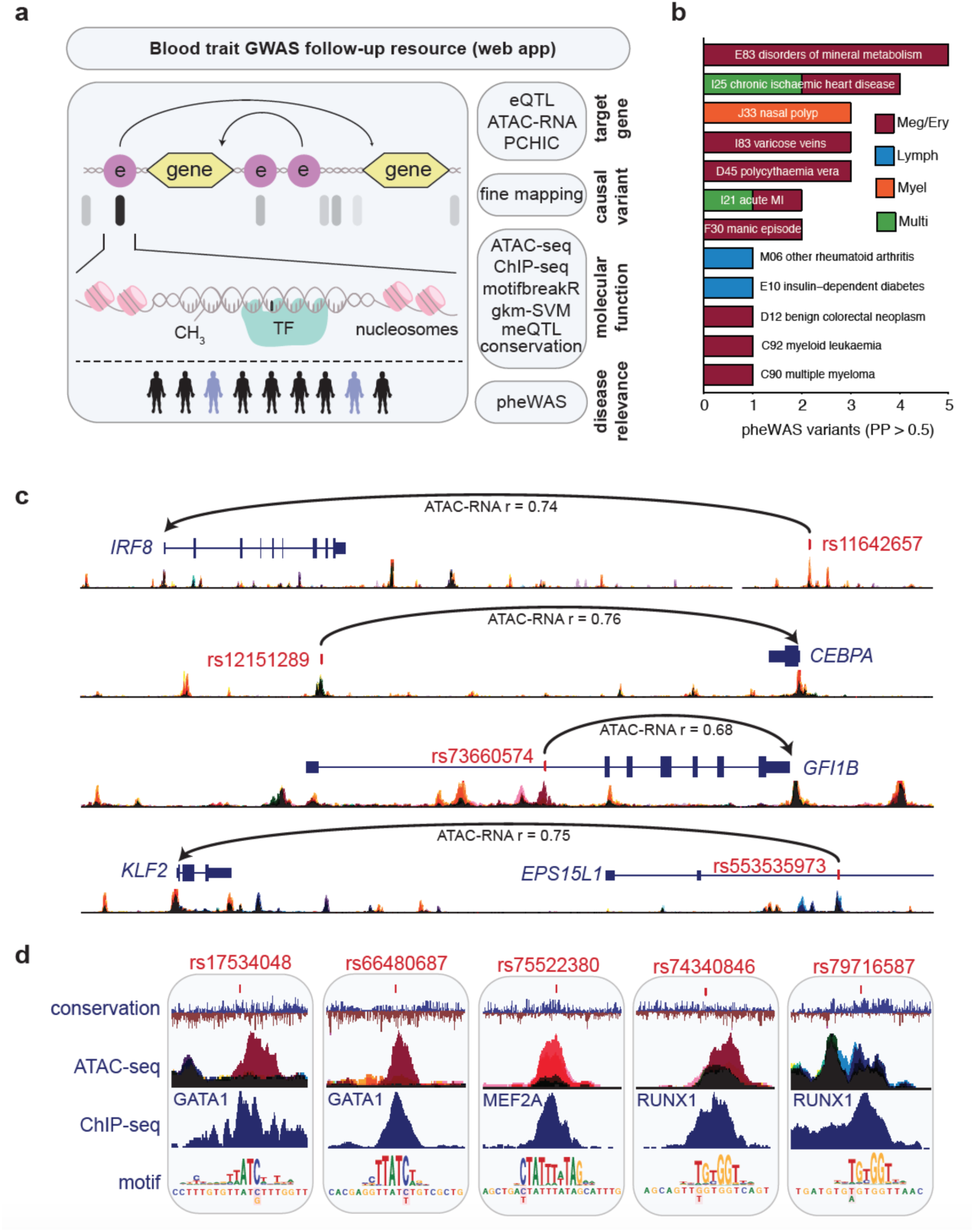
Mechanisms of core gene regulation in blood production, (**a**) Schematic of auxiliary data integrated to predict target genes, mechanisms, and disease relevance for fine-mapped variants. All data are available for use on an interactive web application. (**b**) Top ICD10 codes nominated by UKBB pheWAS analysis for fine-mapped variants (PP > 0.5). Variants identified for each pheWAS trait passed Bonferroni-correction for multiple testing and are annotated based on the lineage of the fine-mapped trait. (**c**) Combining genetic fine-mapping for blood cell traits with dense RNA-seq and ATAC-seq profiling of the hematopoietic tree reveals putative causal variants and their target *trans-acting* TFs (rs11642657 and rs12151289 are associated with platelet traits, rs73660574 is associated with red blood cell traits, and rs553535973 is associated with lymphocyte count). Target genes were identified using ATAC-RNA correlations, and many are additionally supported by promoter-capture Hi-C. Both MEF2A and MEF2C occupied the element in the 3rd panel. (**d**) Additionally combining ChIP-seq with conservation (PhyloP) and motif predictions reveals putative causal variants that disrupt c/s-binding of hematopoietic TFs known to be involved in regulating hematopoietic differentiation for various blood cell traits (rs17534048 and rs66480687 are associated with red blood cell traits, rs75522380 and rs74340846 are associated with platelet traits, and rs79716587 is associated with lymphocyte count; black color represents accessibility throughout hematopoiesis).

To elucidate putative regulatory consequences of our fine-mapped variants, we sought to identify high confidence trans-acting factors that were modulated by GWAS regulatory variants. We identified that *IRF8* and *CEBPA*, two TFs involved in monocyte specification and differentiation,^40,41^ are targets of fine-mapped monocyte count associated variants (**Fig. 6C**). Both variants are within NDRs that are specific to monocyte precursor cells. Similarly, we determined that *GFI1B*, *KLF2*, and *MEF2C* were targets of fine-mapped variants for mean reticulocyte volume, lymphocyte count, and platelet count, respectively, each within a similar progenitor-specific NDR (**Fig. 6C**, **Fig. S11**). In some cases, such as for *IKZF1*, which is broadly expressed in hematopoiesis, several fine-mapped variants for different blood cell traits were determined to interact with the *IKZF1* promoter in a cell type-specific manner (an MCV-associated variant was within an erythroid-specific loop, while a lymphocyte count-associated variant was within a B and T cell-specific loop), consistent with a previous eQTL study of this locus (**Fig. S11**).^42^

Next, we attempted to identify precise cis-regulatory mechanisms of fine-mapped variants. Restricting our hypotheses to the overlap of both biochemical (ChIP-seq) and statistical (canonical motif breaking) results, we posit mechanisms of fine-mapped variants that putatively disrupt TF occupancy in hematopoietic tissues. For example, **Fig. 6D** and **Fig. S11** highlight three fine-mapped variants associated with RBC traits that were within erythroid-specific NDRs, bound by erythroid master TF GATA1, and predicted to disrupt its short DNA binding motif (AGATAA), similar to previously reported functional variants that we had identified proximal to this TF’s binding motif.^43^ We also found fine-mapped variants that were predicted to disrupt RUNX1 binding that were associated with both platelet count^44^ and lymph count,^45^ reflecting the diverse roles of RUNX1 in hematopoiesis (**Fig. 6D**). Interestingly, we were able to identify fine-mapped variants that were predicted to disrupt IRF and MEF2 protein binding in *cis* (**Fig. 6D**, **Fig. S11**). These variants were associated with the same traits as the variants targeting the trans-acting factors (*IRF8* and *MEF2C*), suggesting that *cis-* and *trans-* effects of GWAS may converge as sample size, experimental data, and predictive models evolve and improve.^4^ Given the resolution of our fine-mapped variants, the comprehensiveness of hematopoietic epigenomic measurements, and the breadth of orthogonal measures of function, we believe that our resource will expedite investigation into how these and other fine-mapped variants regulate hematopoietic lineage specification and differentiation.

## Discussion

Two outstanding challenges in the post-GWAS era are **(1)** the precise identification of causal variants within associated loci and **(2)** determining the exact mechanisms by which these variants result in the observed phenotypes, starting with identification of the pertinent cell types. To address **(1)**, we used robust genetic fine-mapping to identify hundreds of putative causal variants for 16 blood cell traits, allowing for up to 5 causal variants in each locus. We combined our fine-mapping results with high resolution chromatin accessibility data for 18 primary hematopoietic populations and derived functional annotations to identify putative target genes, regulatory mechanisms, and disease relevance. Moreover, we highlight several compelling anecdotes for the utility of our approach and provide our fully integrated framework as a community resource.

To address **(2)**, we developed a novel enrichment method (g-chromVAR) that can discriminate between closely related cell types and score single cells for GWAS enrichment, an approach complementary to heritability enrichment methods, such as LDSR. Our new approach allowed us to directly probe the regulatory dynamics of hematopoiesis. Specifically, several recent studies have used various single cell assays to suggest that lineage commitment in hematopoiesis is more complex and occurs earlier than has been classically considered.^17–46^ Here, we used direct phenotypic measurements of blood production at a population level to show that this regulation likely does not occur homogeneously within classically defined populations and is capable of being regulated at the single-cell level by common genetic variation.

Our integrated approach is designed to distinguish closely-related cell types using quantitative chromatin data and reveal specific populations in which trait-associated regulatory variation acts. We suggest that using a well-powered method to identify single cells and cell populations that are trait-relevant provides a critical first step in broadly deciphering causal mechanisms underlying phenotypic variation. Our combined approach further facilitates the identification of *cis-* and *trans-* regulatory programs that are modulated by common genetic variation. As relevant primary tissues and single-cell assays become increasingly available and affordable, we believe that our overall approach will similarly allow for the interrogation of other complex traits at the single-variant and single-cell level.

## Code availability

g-chromVAR is available as an open source R-package distributed freely at http://caleblareau.github.io/gchromVAR. All code required for reproducing results discussed herein is made available at http://github.com/caleblareau/singlecell_bloodtraits.

## URLs

A UCSC Genome Browser visualization hub for all bulk ATAC data is available with this hub URL: https://s3.amazonaws.com/atachematopoesis/hub.txt. The web app to visualize putative causal variants and corresponding annotations is available at http://molpath.shinyapps.io/ShinyHeme.

## Acknowledgements

We want to thank members of the Buenrostro and Sankaran labs for their helpful discussions. This work was supported by National Institutes of Health Grants R01 DK103794 and R33 HL120791 (to V.G.S.) and by the Harvard Society and Broad Institute Fellows programs (to J.D.B.).

## Online Methods

### Genome-wide association studies

Genome-wide association studies were carried out for 16 different blood cell indices in 114,910–116,667 “white British” individuals from UK Biobank. Imputation was performed using the combined 1000 Genomes Phase 3-UK10K panel (http://biobank.ctsu.ox.ac.uk/crystal/refer.cgi?id=157020). To account for population substructure in blood cell traits, we regressed each phenotype against the first 10 principal components, age, and sex. We then inverse normalized the residuals, which were used as the phenotype measurements for the genetic association tests. Specifically, we regressed each phenotype measurement against the probabilistic imputed allele dosage using a linear mixed model approach as implemented in BOLT-LMM v2.2.^47^ Genome-wide significance was defined as a p-value < 5 × 10^−8^.

### Fine-mapping

Sentinel association regions were constructed as follows: first, all variants were ranked by decreasing *χ*^2^ statistics. Next, we derived 3MB regions centered at the top variant - each region is ~3 cMs, so all relevant LD structure should be fully captured for nearly every region (Yu *et al.*^48^ reported that 95% of region recombination rates fall within 3MB). This process was repeated for each top association variant that did not overlap any 3MB regions created thus far until there were no genome-wide significant variants remaining in undefined regions. Within each region, we identified all imputed variants with MAF > 0.1% and imputation quality > 0.6 and calculated z-scores from the summary statistics for each. We next derived dosage LD matrices for each region using LDstore^21^ on the genotype probability files (.bgen) used for the association studies. To be exact, we computed LD matrices from 120,086 individuals who had a phenotype for at least one of the 16 blood cell traits.

Fine-mapping was performed on genome-wide significant GWAS regions using the FINEMAP software with the z-score and LD matrices as input.^22^ The output from FINEMAP is **(1)** a list of potential causal configurations together with their posterior probabilities and Bayes Factors **(2)** the posterior probability marginalized over the causal configurations that individual variants are causal, and **(3)** the posterior probabilities that there are a specific number (between 1 and 5) of statistically independent associations in each region. Default FINEMAP settings were used and we kept all variants with posterior probabilities > 0.1% for downstream analyses. For the *CCND3* and *AK3* regions in which follow-up luciferase reporters were performed, we reran FINEMAP allowing for up to 10 causal variants, confirming ~4 independent effects in the *CCND3* locus (60.6% posterior probability) but revealing ~8 independent effects for the *AK3* locus (59.9% posterior probability).

To confirm select regions with multiple putative causal variants, we performed conditional analysis using BOLT-LMM by first conditioning on the variant with the lowest p-value in the region and then stepwise adding to the model the variant with the lowest conditional p-value until no additional variant reached the genome-wide significance threshold of 5×10^−8^ in the combined model.

### ATAC and scATAC sequencing and data preprocessing

Chromatin accessibility profiles for a total of 18 populations, including 15 previously reported,^27,28^ were assayed using FastATAC, an optimized ATAC-seq protocol optimized for primary blood cells, as previously described.^28,49^ Sequencing data for each of the 18 populations was uniformly processed using a custom pipeline that includes sequencing adaptor removal, alignment using Bowtie2,^50^ and PCR duplicate removal with Picard RemoveDups command.

Accessible chromatin peaks were called from the 18 sorted populations of blood cells using MACS2.^51^ To derive a consensus set of loci for downstream analysis, individual peaks were resized to a uniform width of 500 bp, centered at the summit from the MACS2 call as previously described.^28^ To derive a consensus peak set for the blood cell types, peaks were combined by removing any other peak overlapping with a peak with greater signal at the summit within a particular cell type. A total of 451,283 peaks representing a consensus set across these 18 sorted bulk populations were called. The average number of fragments in this consensus peak set ranged from 4.4 million (pDCs) to 37.1 million (CMPs) for a mean of 19.3 million reads in peaks per sorted cell type (**Table S3**).

FACS-sorted cells from 9 distinct cellular populations from CD34+ human bone marrow, which included cell types spanning the myeloid, erythroid, and lymphoid lineages, were additionally profiled as previously described.^27,28^ Single-cells were sorted then assayed using scATAC-seq^27,52^ across a total of 30 independent single-cell experiments representing 6 human donors, with each population assayed from two or more distinct donors. In total, our raw data set comprised 3,072 single-cell chromatin accessibility landscapes with 2,034 cells passing stringent quality filtering. These cells yielding a median of 8,268 fragments per cell with 76% of those fragments mapping to peaks, resulting in a median of 6,442 fragments in peaks per cell again using a consensus peak set that was inferred for these specific progenitor populations.^27^

To infer dynamic GWAS enrichments across hematopoietic differentiation, pseudotime orderings of single cells across three lineages (erythroid, lymphoid, and myeloid) were estimated using an adaptation of the Waterfall algorithm^53^ as previously described.^27^ To assess regulatory heterogeneity of single cells, we computed a chi-squared statistic for each trait/cell type’s z-scores to test whether the observed variance was greater than expected. Under the null, the variance of z-scores is 1 from the definition of our statistic (see *g-chromVAR* methods below), and we observed greater variation than expected only for traits within the CMP and MEP populations. Within CMP and MEP populations, we applied k-medoids clustering on the first 5 principal components within each sorted population from global chromatin accessibility profiles for each cell.^27^ For both the CMPs and MEPs, the optimal cluster number was determined by maximum average silhouette width. Post-hoc analyses of heterogeneity within the partitioned clusters of the erythroid-enriched CMPs confirmed that megakaryocyte-erythroid enrichment was not distinct within CMPs.

### g-chromVAR

The bias-corrected enrichment statistics for ***T*** traits and a set of ***S*** samples (chromatin cell type profiles) with ***P*** peaks computed by g-chromVAR is a generalization of the chromVAR method.^26^ Intuitively, our implementation of g-chromVAR relaxes the requirement in chromVAR that trait-peak annotations be binary, allowing for uncertainty in annotations such as transcription factor binding or in our case, localization of GWAS variants. Specifically, the chromVAR implementation requires a binarized matrix ***M*** (dimension ***P*** by ***S***) where *m*_*i*,*k*_is 1 if annotation *k* is present in peak *i* and 0 otherwise. For example, in our examination of chromVAR (**Figure 3b,d**), ***M*** represents a binary matrix where *m*_*i*,*k*_ = 1 if a genome-wide significant variant for trait *k* was present in peak *i*. However, our application of chromVAR to variant association data for our 16 hematopoietic traits revealed inflated summary statistics and poor lineage-specific enrichments without modeling the posterior confidence of variants (**Figure 3a**). We note that if FINEMAP identified only 1 causal variant per region with a posterior probability of 1, g-chromVAR and chromVAR would yield identical results.

Instead, our methodology, g-chromVAR, uses a matrix of variant posterior probabilities ***G***, where *g*_*i*,*k*_ is the sum of the posterior probabilities of the variants contained in the genomic coordinates of peak *i* for each trait *k*. Using the matrix of fragment counts in peaks ***X***, where ***x***_*i*,*j*_ represents the number of fragments from peak *i* in sample *j*, a matrix multiplication ***X***^*T*^ · ***G*** yields the total number of fragments weighted by the fine-mapped variant posterior probabilities for *S* samples (rows) and ***T*** traits (columns). To compute a raw weighted accessibility deviation, we compute the expected number of fragments per peak per sample in ***E***, where *e*_*i*,*j*_ is computed as the proportion of all fragments across all samples mapping to the specific peak multiplied by the total number of fragments in peaks for that sample:

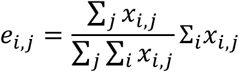

Analogously, ***X***^***T***^ · ***E*** yields the expected number of fragments weighted by the fine-mapped variant posterior probabilities for ***S*** samples (rows) and ***T*** traits (columns). Using the ***G***, ***X***, and ***E*** matrices, we then compute the raw weighted accessibility deviation matrix ***y*** for each sample *j* and trait *k* (*y*_*j*, *k*_) as follows:

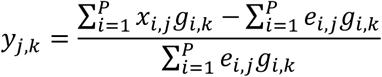

To correct for technical confounders present in assays (differential PCR amplification or variable Tn5 tagmentation conditions), g-chromVAR borrows the strategy implemented in chromVAR by generating a background set of peaks intrinsic to the set of epigenetic data examined. We note that other GWAS enrichment tools such as LDSR or GoShifter ignore biases prevalent in epigenomic assays that are explicitly corrected by g-chromVAR. In particular, variance in PCR or Tn5 tagmentation quality can lead to substantial differences in the number of observed fragment counts between cells based on an individual peak’s GC content or average accessibility,^26^ leading to errant GWAS-cell type enrichments. To correct for these technical confounders, each peak is assigned a background set of peaks that are matched in mean nucleotide GC content and average fragment accessibility between the sums of the cell types. An inverse Cholesky transformation is applied to a *P* by 2 matrix containing these variables to generate two uncorrelated dimensions describing the per-peak confounding. This two-dimensional space is divided into a pre-defined number of equally spaced bins where bin *i* is indicated *β*_*i*_. Each peak *q* is assigned a bin from the shortest Euclidean distance between the bin’s centroid and the individual peak in this transformed space. The probability that a peak *q*′ in bin *j* is selected as a background peak for peak q is proportional to the distance between bins *i* and *j* over the total number of peaks in bin ***j*** |*β*_*j*_|:

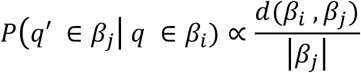

where the distance function *d* contains hyperparameters which, along with the total number of bins, have been previously discussed.^26^

By default, the framework samples a background set of 50 background elements per peak, which we’ve verified to be robust (**Fig. S4B**). The matrix ***B***^(***b***)^ encodes this background peak mapping where 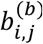 is 1 if peak *i* has peak *j* as its background peak in the *b* background set (*b* ∈ {1,2, …,50}) and 0 otherwise. The matrices ***B***^(***b***)^ · ***X*** and ***B***^(***b***)^ · ***E*** thus give an intermediate for the observed and expected counts also of dimension ***P*** by ***S***. For each background set *b*, βsample *j*, and trait *k*, the elements 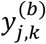 of the background weighted accessibility deviations matrix ***y***^(*b*)^ are computed as follows:

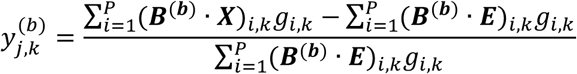

After the background deviations are computed over the 50 sets, the bias-corrected matrix ***Z*** for sample *j* and trait *k* (*z*_*j*,*k*_) can be computed as follows:

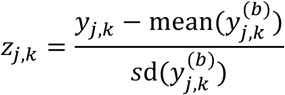

where the mean and variance of 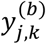 is taken over all values of *b* (*b* ∈ {1,2, …,50}). Sample-trait p-values can then be computed from the one-tailed normal distribution of these z-scores using the pnorm function in R. Our implementation of g-chromVAR utilizes efficient matrix operations for each step and can compute pair-wise trait-cell type enrichments in ~1 minute on a standard laptop computer.

### Simulations

To verify that enrichments computed by g-chromVAR were well-calibrated in our system, we devised a general simulation framework that computes enrichments for the 18 bulk hematopoietic cell types for an arbitrary simulated phenotype. Using the same matrix of fragment counts in peaks ***X***, where *x*_*i*,*j*_ represents the number of fragments from peak *i* in sample *j*, we simulated a causal relationship between one of the accessibility samples *j* by performing a weighted draw of observed variant posterior probabilities ***G***, where *g*_*i*,*k*_ is the sum of the posterior probabilities of the variants contained in the genomic coordinates of peak *i* for each trait *k*.

Specially, we first perform a counts-per-million (CPM) transformation of the fragment counts in peaks matrix to account for uneven sequencing depth between samples. Next, we z-transform the CPM-normalized matrix row wise to yield a matrix termed ***X***\* where 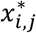 represents the amount of open chromatin observed in sample *j* in peak *i* relative to other samples. Intuitively, elements of the z-score matrix ***X***\* yield larger positive numbers for cell type-specific peaks in specific samples (values 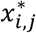 range from −3.46 to 4.01). This matrix ***X***\* serves as a basis for determining the celltype specificity of an individual regulatory element.

To generate simulated elements of ***G***, we define a sorted vector *ν* of length ***T*** * ***P*** (where 99.7% of values were zero) from the observed elements of ***G*** for our ***T*** = 16 hematopoietic traits and ***P*** = 451,283 regulatory peak elements. This vector *ν* thus represents empirically derived values from the hematopoietic system studied that serve as input into g-chromVAR. Then, for a fixed causal cell-type *j*, we generate matrix] of q ∈ (1,2,…, 100) simulated traits, where entries are defined as follows:

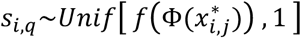

Here, *f* is a linear function that maps the normal cumulative distribution function (CDF) transformation of the 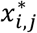 z-score to a (0, 1) real number and is calibrated to yield phenotypic values similar to those observed empirically (matched mean column sum of ***G***). The *s*_*i*,*q*_ value thus is a randomly generated (0, 1) real number skewed toward 1 when peak *i* contains cell type-specific chromatin for the fixed cell-type *j*. A final transformation of ***S*** maps these (0, 1) real number values to observed weights (elements of ***G*** or equivalently *ν*) using the inverse CDF of *ν* to index values. This final matrix, which serves as the input for g-chromVAR, is simulated to be enriched for cell-type *j* and null for all others. For the fully null simulation, elements of *s*_*i*,*q*_ were populated from random draws of a *Unif*[ 0,1].

### Linkage disequilibrium score regression

We used LD score regression (LDSR) to compute the narrow-sense heritability estimates and genetic correlations of the 16 blood cell traits in the UKBB. Reference LD scores were computed with a subset of unrelated European individuals from the UK10K cohort. To remove genetically related individuals, we first used Plink to construct a filtered list of variants with MAF > 0.10 and no pair of variants with R^2^ > 0.10. These LD and MAF-pruned variants were then used to calculate an identity-by-descent (IBD) matrix, and one individual in each pair of samples with proportion IBD (*π̂*) > 0.125 were removed to produce a final subset of 3,677 unrelated individuals to serve as the reference panel for LDSR. After applying the recommended variant filtering, z-scores for an average of 6,655,000 variants per trait were used as input to LDSR. For heritability estimates for variants identified by fine-mapping or by linkage to the sentinel, we note that these estimates may be either over-estimated or under-estimated from the reported values as previously noted.^54^

To estimate cell type enrichment for each trait, stratified LDSR was used to partition each trait’s heritability into the baseline model of 53 annotations, as well as each of the 18 hematopoietic ATAC-seq annotations (one at a time). P-values for cell-type enrichment were derived from the z-scores of the coefficients for each hematopoietic annotation with statistical significance assessed at a stringent Bonferroni threshold of 0.00017 (corrected for 16 traits and 18 cell types).

### Local Annotation Shifting

We implemented a slightly modified version of GoShifter to calculate the enrichment between fine-mapped variants with PP>0.01 for every trait and 5 different genomic annotations. To obtain the annotation for hematopoietic nucleosome-depleted regions (NDRs), we used the consensus peak set for all blood cell types, performed row and column quantile normalization on the counts matrix, and kept only peaks that had a maximum count in the top 80% for at least one of the 18 cell types. The coding, intron, promoter, and 5' untranslated region annotations were obtained from the UCSC Genome Browser as previously processed.^55^

To calculate enrichments, we calculated the overlap between the fine-mapped variant set of each trait (16 total) with each of the 5 genomic annotations. To define the null distribution of annotation overlap, we performed 10,000 locally shifting permutations; with every permutation, we shifted the genomic coordinates of the fine-mapped variant set by a random distance between −1.5MB and 1.5MB (this approach is equivalent to shifting the annotations). This was performed using the permTest function of the regioneR package. The final odds ratio was calculated by dividing the number of overlaps between the original fine-mapped variant set and a genomic annotation by the mean number of overlaps between the 10,000 permuted sets and the same genomic annotation. To test if the association between a fine-mapped variant set and a genomic annotation (e.g. hematopoietic NDRs) was highly dependent on their exact position, we used the localZScore function to calculate enrichment scores after various increments of shifting the fine-mapped variant set.

### Bulk RNA-seq samples and peak-gene correlations

Raw sequencing reads from sorted populations from bulk RNA-seq experiments previously described^27,28^ were aligned to the hg19 reference genome using STAR version 2.5.1b^56^ with default parameters. Per-gene transcript quantifications were summed over biological and technical replicates to provide a single transcript count per sorted cell type for 17 total populations matching the analogous bulk ATAC profiles (RNA for megakaryocytes was absent). To determine empirical peak-gene associations, Pearson correlation was computed for each peak within a 1MB window of the transcription start site per gene using the log counts per million value for each feature.

### Luciferase Reporter Analysis

Firefly luciferase reporter constructs (pGL4.24) were generated by cloning the variant(s) of interest centered in 300–400 nucleotides (*AK3* 325bp; *CCND3* 363bp) of genomic context upstream of the minimal promoter using BglII and XhoI sites. The Firefly constructs (500ng) were co-transfected with a pRL-SV40 Renilla luciferase construct (50ng) into 100,000 K562 cells using Lipofectamine LTX (Invitrogen) according to manufacturer’s protocol. After 48 hours, luciferase activity was measured by Dual-Glo Luciferase assay system (Promega) according to manufacturer’s protocol. For each sample, the ratio of Firefly to Renilla luminescence was measured and normalized to the empty pGL4.24 construct.

A total of four haplotypes were constructed per locus to examine the effects of two fine-mapped putative causal variants. For the *CCND3* locus, we examined the effects of rs112233623 (ref: C, alt: T) and rs9349205 (ref: **G**, alt: A), which are 161 base pairs apart. For the *AK3* locus, we examined rs409950 (ref: A; alt: C) and rs12005199 (ref: A, alt: G), which are separated by 123 base pairs. A total of nine (n = 9) experimental replicates per haplotype (four haplotypes per locus), including the empty pGL construct, were measured across two experimental batches.

To compute the additive and multiplicative effects of each variant, we used a generalized linear model of the following form for both of the *AK3* and *CCND3* loci separately:

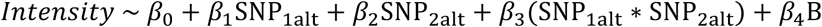

Here, the luciferase intensity is defined as the ratio of Firefly to Renilla luminescence normalized to the empty vector for each experimental replicate. The additive effects of the two SNPs were estimated using *β*_1_ and *β*_2_ whereas the multiplicative effect of both SNPs on the same haplotype was computed using an interaction term, *β*_3_. We encoded each variable such that the reference allele was a 0 whereas the alternate allele was a 1 for each experimental sample. Finally, we adjusted for variable infection efficiency between the experimental batches using a fixed effect variable *B* (*B* ∈ {0,1}). To increase power, point estimates and standard errors were realized directly from the linear model using the *β* coefficients from each reporter set rather than the mean of the specific haplotype.

### Transcription factor motif analysis

Prediction of the effects of fine-mapped variants on transcription factor binding sites (TFBS) was performed using the motifbreakR package^36^ and a comprehensive collection of human TFBS models (HOCOMOCO^35^). All fine-mapped variants with PP>0.1%, a singlenucleotide substitution, and a dbSNP rsID were supplied as input. We applied the “information content” scoring algorithm and used a p-value cutoff at 5×10^‒4^ for a TFBS match; all other parameters were kept at default settings.

### PheWAS

To identify potential disease roles for our fine-mapped variants, we performed a phenome-wide association study (pheWAS) for each variant using publicly available summary statistics from the UK BioBank (http://www.nealelab.is/blog/2017/7/19/rapid-gwas-of-thousands-of-phenotypes-for-337000-samples-in-the-uk-biobank). We investigated 642 unique International Statistical Classification of Diseases and Related Health Problems, 10th Revision (ICD-10) medical billing codes. Each individual with a code was treated as a case and all remaining individuals were treated as controls. We investigated whether 729 distinct variants with a FINEMAP PP > 50% were also associated with these 642 medical diagnoses, resulting in 468,018 (729*642) tests, leading to a Bonferroni-adjusted threshold of 1.07×10^−7^.

### gkm-SVM

We used gkm-SVM^38^ to predict the effects of our fine-mapped variants on open chromatin *in silico*. Specifically, we selected ATAC peaks (peaks with counts > 90th percentile) from each of the 18 cell types (~ 50,000 to 60,000 for each cell type) as positive sequence sets. We generated 81,763 negative control peaks matching the GC, length, and repeat content of the full peak set. We then trained gkm-SVMs for each cell type using 11-mers and 5-fold cross validation (average area under the precision recall curve was 93%). For each variant, we calculated a score using the deltaSVM method, where we sum all possible spanning 11-mers for the alternate allele and subtract the same sum from the reference allele.

### eQTL analysis

To identify putative gene regulatory targets of our fine-mapped variants, we intersected variants with a PP > 0.001 with previously reported variant-gene associations (P < 5e-8) derived from whole-blood eQTL (both *cis* and *trans*) associations from two previous studies of 5,311^42^ and 2,765^57^ individuals. In addition, we intersected fine-mapped variants with previously reported variant-CpG methylation status associations (P < 5e-8) from a recent study of 1,366 individuals^58^.Data was processed by SMR and downloaded from http://cnsgenomics.com/software/smr/.

